# Label-Free Mapping of Subcellular Dynamics using Wide-field Interferometric Scattering Microscopy and Spectral Exponent Analysis

**DOI:** 10.1101/2025.06.23.661093

**Authors:** Caroline Livan Anyi, Hengze You, Huakun Li, Kheng Ling Goh, Tong Ling

## Abstract

Interferometric scattering (iSCAT) microscopy combines label-free detection with nanometer-scale motion sensitivity. It captures temporal signal fluctuations from sub-diffraction movements of scatterers in live cells, including macromolecules and cell membranes. Here, we investigate label-free mapping of subcellular dynamics by calculating the power spectral density (PSD) of the iSCAT time series. Using a wide-field iSCAT, we observe an inverse-power-law relationship *S*(*f*) = *βf*^−*α*^ over 30 - 1250 Hz in multiple cell types. We compute pixel-wise PSDs and obtain a color-coded spectral exponent map based on the fitted spectral exponent (*α*) and amplitude (*β*), which indicates the characteristics and strength of subcellular movements. We also incorporate the goodness-of-fit as color saturation to enhance the contrast of the spectral exponent map. Following this approach, we demonstrate label-free imaging of live cells while effectively distinguishing between benign and malignant thyroid cancer subtypes, mitotic and interphase cells, as well as live and apoptotic cells. Dynamic imaging using wide-field iSCAT provides an intrinsic, label-free marker of cellular states and mechanical properties, which may benefit studies in mechanobiology, cancer diagnostics, and stem cell therapies.

## 1. Introduction

Biological cells are highly heterogeneous structures composed of numerous internal compartments, each performing specific functions. These compartments (such as the cell nucleus, endoplasmic reticulum, and mitochondria) feature unique structures and intricate topologies that lead to distinct thermal motion at the submicron to nanometer scale (*1–3*). Statistical analysis of these random thermal motions can infer the size of the object and its interaction with the surrounding environment (*4–9*).

Light scattering-based imaging techniques, which utilize light interaction with subcellular structures, provide a label-free and non-invasive approach to studying live cells. One such powerful imaging modality is interferometric scattering (iSCAT) microscopy. iSCAT microscopy uses common-path interferometry to detect elastic scattering fields, offering exceptional sensitivity for label-free imaging of nano-objects (*10–13*). Regarding live cell imaging, Park et al. utilized iSCAT to image live mammalian cells on the coverslip, revealing interference fringes over the lamellipodia caused by reflections from the plasma membrane and the cover glass interface (*14*). To overcome unwanted reflections, Hsieh and colleagues proposed coherent brightfield microscopy (COBRI) that detects forward-scattering signals and utilizes non-scattered transmitted light as the reference beam (*15*). Using COBRI, Huang et al. resolved the 3D motion of single virus particles and intracellular vesicles in live cells with nanometer precision and a temporal resolution of 10 μs (*16*, *17*).

Despite the above advantages, imaging live subcellular structures using a wide-field iSCAT is often challenging due to the speckle patterns arising from the self-interference of light scattered from the various scatterers in live cells (*18*, *19*). These speckle patterns produce intensity fluctuations that complicate high-resolution imaging and signal analysis. On the other hand, the statistical information of their fluctuations in time infers cellular activities and microscopic mechanical constraints, which have been actively investigated in biomedical imaging. For example, dynamic optical coherence tomography (OCT) leverages the temporal fluctuations in the OCT signal for label-free imaging of thick tissues and organoids (*20*, *21*). The proposed analytic techniques for dynamic OCT include calculating the mean frequency of the PSD (*22*, *23*), logarithmic intensity variance-based metrics and OCT-correlation-decay-speed (*24*, *25*). In iSCAT, Hsiao et al. analyzed speckle fluctuations in live cell nuclei (*8*, *26*). By detecting light scattered from unlabeled chromatin, they captured molecular diffusion and active processes within the nucleus. Chromatin movement resulted in time-varying speckle patterns, with short-term fluctuations on the millisecond timescale characterized by variance strongly correlating with local chromatin density (*26*). Label-free, chromatin-specific imaging can thus be achieved by extracting high-frequency temporal signal fluctuations and analyzing their statistics.

Stochastic movements at the microscopic scale driven by thermal energy, such as Brownian motion, molecular diffusion and membrane fluctuation, often result in a scale-invariant power-law relationship when calculating the PSD of the sample movement over time (*27–29*). In this paper, we investigate whether a similar phenomenon can be observed in an ensemble manner in label-free iSCAT imaging of live cells. Specifically, we evaluate whether the PSDs of the iSCAT signals in live cells conform to a power-law relationship expressed as *S*(*f*) = *βf*^−*α*^, where *α* represents the spectral exponent and *β* represents the amplitude. We construct a 2-D color-coded spectral exponent map based on the fitted parameters. To demonstrate the applicability of the proposed approach, we validate its utilization across various cell lines, including spiking human embryonic kidney (HEK) cells and tumor cells (U2OS, HeLa, and thyroid carcinoma) under several biologically relevant scenarios: (i) live and fixed cells, (ii) cells undergoing mitosis, (iii) cells undergoing apoptosis, and (iv) variations in malignancy among tumor cells.

## 2. Results

### 2.1 High-speed iSCAT imaging of live cells reveals power-law relationship in the PSD of iSCAT signal variation

The principle of iSCAT imaging is illustrated in Fig. 1(a). In our experiment of live cell imaging, the incident light *E*_inc_ was focused onto the back focal plane of an objective lens via a lens, which generated a wide-field illumination on the sample, and the reference light *E*_ref_ was produced by the reflection from the top surface of the glass coverslip due to the refractive index difference between the sample medium and glass interface. Both the reference light *E*_ref_ and the light scattered from scatterers within the sample *E*_sca_were collected by the objective lens and directed to the imaging path via a beamsplitter. The interference between *E*_ref_ and *E*_sca_ resulted in the iSCAT image series over time, as shown in Fig. 1(b).

**Figure 1:**
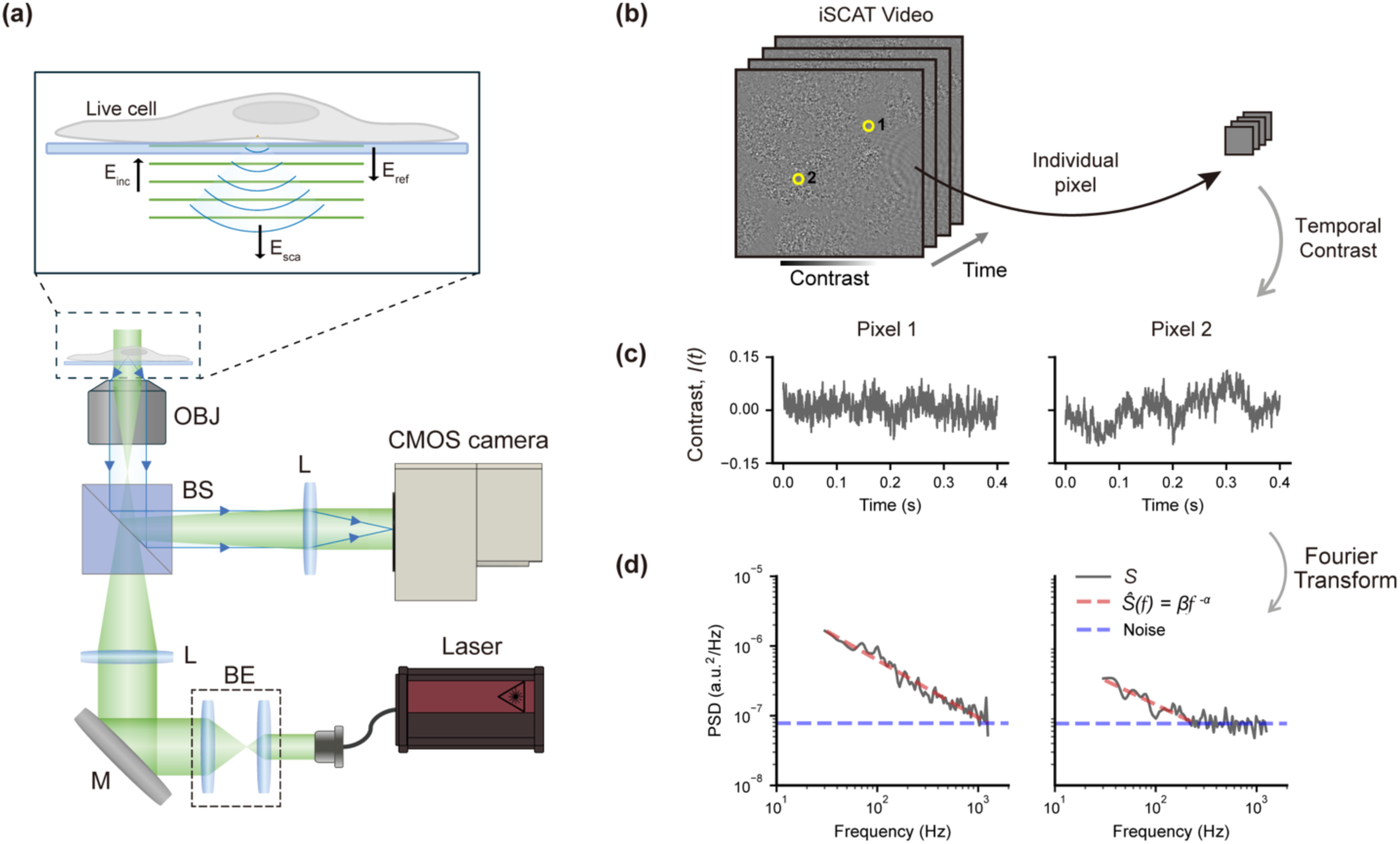
Interferometric scattering imaging and power spectral analysis on live cells. (a) Schematic illustration of the iSCAT imaging concept for live cells. OBJ: Objective lens, BS: Beam splitter, BE: Beam expander, L: Achromatic lens, M: Silver mirror. (b) An example of an iSCAT contrast video. The yellow circles indicate two representative pixels analyzed in the subsequent panels. (c) Temporal contrast fluctuations of the two pixels over time. (d) PSD analysis of the corresponding temporal signals. The PSDs are fitted to Eq. (3), represented by the dashed red lines.

To mitigate the speckle patterns caused by the scatterers’ crosstalk, we used broadband illumination (530– 700 nm) from a supercontinuum laser that reduced the coherence length to ∼0.8 μm. In addition, we employed a wide-field, low numerical aperture (NA) objective (20×, NA=0.75) with a depth of focus of ∼2 μm and intentionally adjusted the focus to the cells’ basal membrane. These configurations effectively reduced the influence of the Gouy phase in the scattered light. If we neglect the contribution of the Gouy phase and assume that the glass-water interface represents the plane with zero optical path length, the detected raw intensity *I*_*d*_(*t*) on the CMOS camera can be expressed as (*30*),

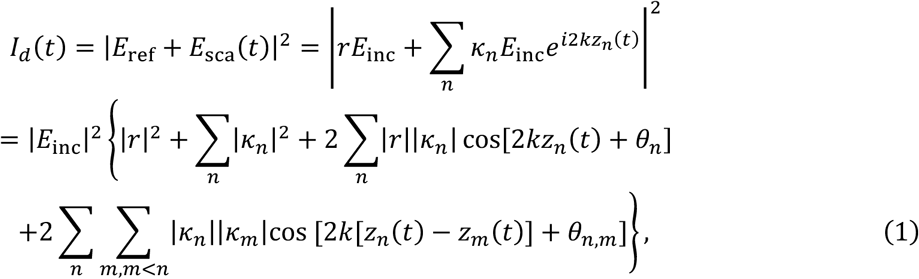

where *r* denotes the reflection coefficient of the glass-water interface, *k* represents the center wave number, *z*_*n*_(*t*) represents the optical path difference between the reference light and the light scattered from the *n*-th scatterer. *κ*_*n*_ denotes the complex polarizabilty of the *n* -th scatterer. *θ*_*n*_ = arg *κ*_*n*_ − arg *r*, which accounts for the phase difference between *r* and *κ*_*n*_, while *θ*_*n*,*m*_ = arg *κ*_*n*_ − arg *κ*_*m*_ represents the phase difference among the complex polarizabilities of scatterer pairs.

Given the significantly weaker magnitude of the complex polarizabilities of each scatterer (membranes, coacervates, macromolecules, etc.) compared to the reflection coefficient of the glass-water interface, along with the reduction of crosstalk among scatterers using a short coherence length (∼0.8 μm), we expect that the majority of our raw iSCAT signal originates from the first and third terms of Eq. (1). Following the derivation outlined in Supplementary Discussion 1, we can obtain the iSCAT contrast signal in our experiments as,

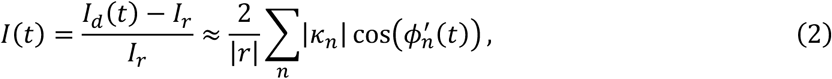

where 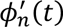 is the overall phase difference between the *n*-th scatterer and the reference light at time *t*.

The contrast signals *I*(*t*) extracted from two representative pixels of an iSCAT video are shown in Fig. 1(c). When we performed the Fourier transform on *I*(*t*) to compute the PSD spectrum *S*(*f*), we observed a PSD that exhibits a power-law relationship within the frequency range of 30 - 1250 Hz (pixel 1). For pixel 2, as the signal amplitude decreased at higher frequencies, this trend faded into a band of white noise, presumably due to shot noise. Similar phenomena were observed in nearly every region occupied by cells, which is further elaborated in the next section.

### 2.2 Spectral exponent analysis of live cell iSCAT signals in full field

We examined whether the power-law relationship observed in Section 2.1 could apply to the iSCAT signals obtained from an arbitrary region of live cells or if it had specific origins. We fitted an inverse-power-law model,

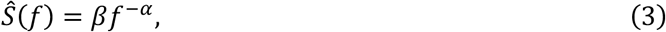

to the PSDs of the iSCAT contrast signals across all pixels, using Welch’s method and curve fitting based on a piecewise function (see Methods and Supplementary Discussion 2). Surprisingly, almost all pixels within the cells exhibited a strong match between the experimental observation *S*(*f*) and the model estimations *Ŝ*(*f*), as evidenced by the high goodness-of-fit values shown in Fig. 2(a).

**Figure 2:**
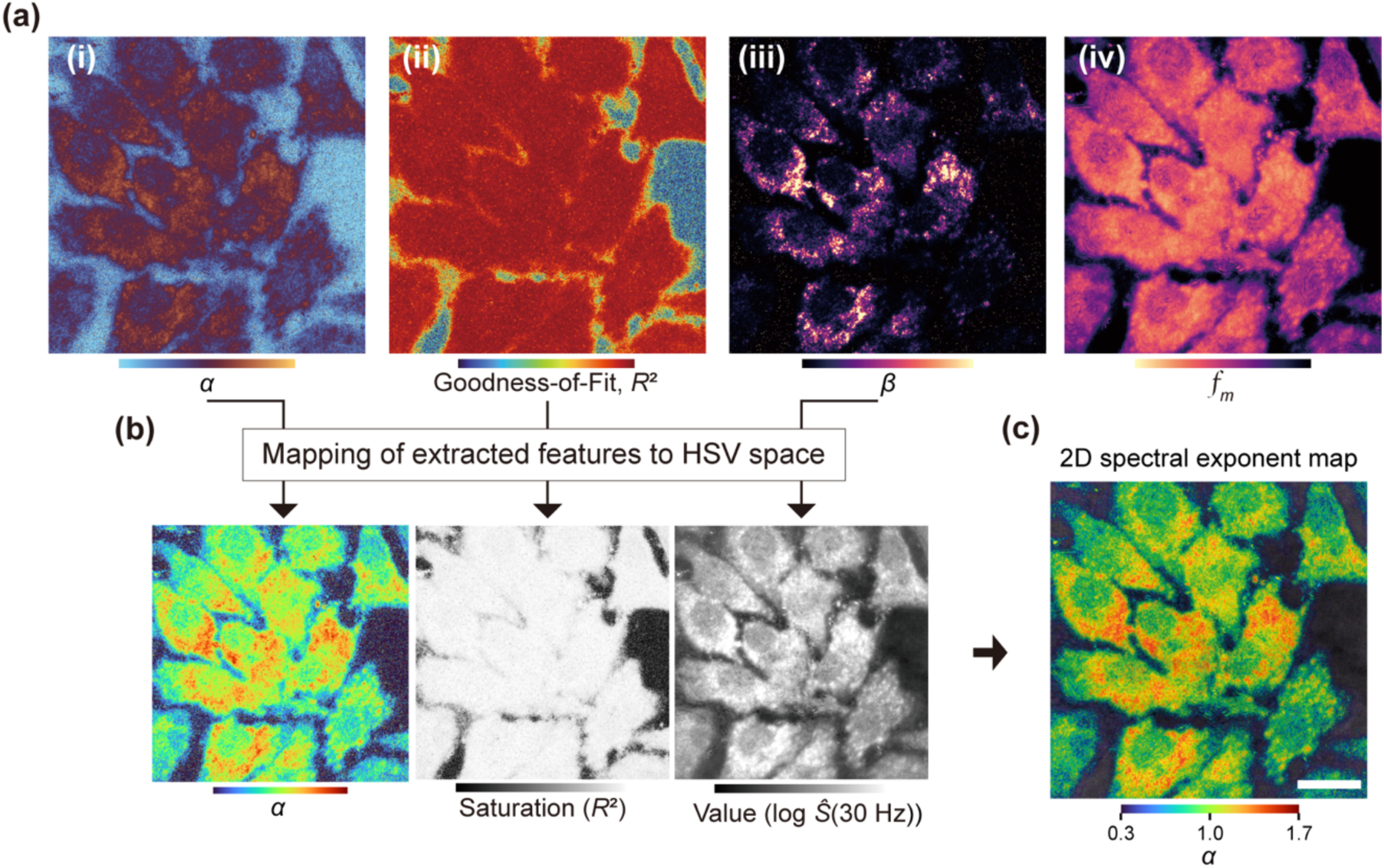
Spectral exponent mapping visualizes the subcellular dynamics of live U2OS cells in full field. (a) Heatmaps of fitted power-law parameters extracted from live U2OS cells: (i) *α*-map, (ii) Goodness-of-fit (*R*^2^) map, (iii) *β*-map, and (iv) *f*_*m*_-map. (b) From left to right: the raw color-encoded *α*-map, the *R*^2^ map fed into the saturation channel in the HSV space, and the amplitude map of *S*&(*f*) value at *f* = 30 Hz fed into the brightness channel in the HSV space. (c) 2D spectral exponent map constructed from (b). Scale bar: 20 µm.

In light of the ubiquity of the power-law relationship observed in live cell imaging, we constructed several spatial maps from power spectral analysis to visualize the dynamic behavior of the sample. The *α*-map in Fig. 2(a) provides insights into how the PSD decays with frequency across the sample. This *α*-map allows us to characterize the underlying nature of the dynamics in different regions. Specifically, regions where the PSD is flat (*α* ≈ 0) indicate that all frequencies contribute equally, a signature of white noise. Considering the model of ensembled iSCAT signals from monodispersed scatterers in solution (see Supplementary Discussion 1), regions with *α* = 2 reflect a typical decay rate for systems dominated by Brownian motion or classical diffusion. An *α* value between 1 and 2 suggests anomalous sub-diffusion in biological systems due to structural constraints or macromolecular crowding, whereas an *α* value larger than 2 indicates super-diffusion resulting from active movements driven by molecular motors or directed transport (*31–33*).

The R²-map in Fig. 2(a)(ii) illustrates the goodness of fit to the power-law model. Meanwhile, the *β***-**map in Fig. 2(a)(iii) indicates the variation in *β* values corresponding to regions with either minimal or significant dynamics. A 2D spectral exponent map (Fig. 2(c)) was created to visualize the spatial distribution of the fitted power-law parameters simultaneously (See Methods). Additionally, we obtained the mean frequency, *f*_*m*_-map (Fig. 2(a)(iv)), which is commonly used for dynamic OCT imaging (*20*), for comparison in the performance. We tested the proposed spectral exponent analysis in both live and fixed U2OS cells to identify the typical spectral parameters associated with active versus inactive states (See Supplementary Discussion 4).

### 2.3 Spectral exponent analysis of mitotic progression in HeLa cells

Time-lapse iSCAT videos were acquired in HeLa cells undergoing mitotic progression. Fig. 3 shows three HeLa cells identified in their early mitotic phase. At the 0-minute mark, these cells displayed a rounded morphology, a typical feature of pro/metaphase, accompanied by relatively high medium *α*-values (∼0.83) and dominant yellow hues in the 2D spectral exponent map. Between 30 to 45 minutes, the cell progressed through anaphase and cytokinesis, forming two daughter cells. The medium *α*-values during these phases decreased to ∼0.7. By 100 minutes, the cells had entered interphase, during which *α* -value further decreased to a level similar to the surrounding non-dividing cells.

**Figure 3:**
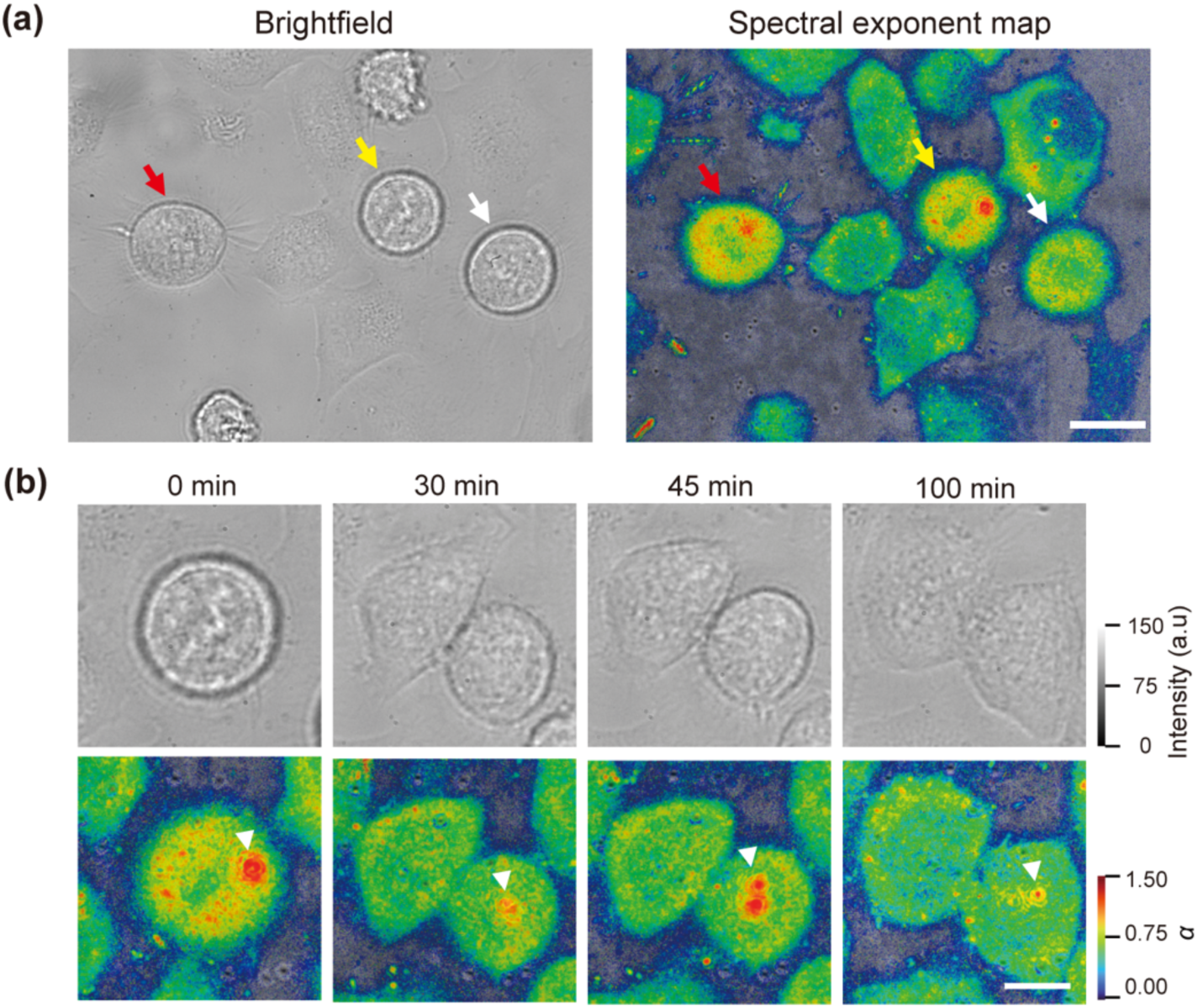
HeLa cells in different phases of mitosis. (a) Brightfield image and 2D spectral exponent map of HeLa Cells. The arrows indicate the cells in the process of mitosis. Scale bar: 20 µm; (b) Time-lapse zoom-in view of the brightfield and 2D spectral exponent maps of the HeLa cell indicated by the yellow arrow in (a) at four time points (0 min, 30 min, 45 min, and 100 min). The white triangular arrows in the 2D spectral exponent maps (bottom row) indicate gold nanoparticles internalized by the cell. Scale bar: 10 µm.

Additionally, we observed two 200 nm gold nanoparticles (AuNPs) internalized by HeLa cells, as indicated by the white triangular arrows. Compared with the intrinsic spectral components of live cells, these AuNPs exhibited significantly higher α-values, likely reflecting distinct biophysics resulting from nanoparticle-cell interactions (Fig. 3(b)). Notably, the same AuNPs cannot be distinguished in the brightfield images, which highlights the advantage of the extra motion contrast provided by the iSCAT-based dynamic imaging.

### 2.4 Identifying live and apoptotic cells in 2D spectral exponent maps

In addition to mitotic progression, we also investigated the dynamic behavior during apoptosis, another cellular process that entails extensive intracellular reorganization. Apoptosis is an irreversible pathway of programmed cell death characterized by distinct fluctuations, particularly at the plasma membrane. We induced apoptosis in spiking HEK cells via hydrogen peroxide (H₂O₂) treatment, which generated oxidative stress that led to membrane damage and cellular degradation (*34*). Over the course of an hour following the H₂O₂ treatment, membrane blebbing was observed in brightfield images, as shown in Fig. 4(a). The 2D spectral exponent maps of those blebbing areas revealed *α* values as high as 1.3. Since blebbing is formed by plasma membrane without additional mechanical support from the actin cortex, the movement of membrane blebbing should be primarily dominated by thermally excited undulations of the lipid bilayer (*35*). This may explain the exceedingly high spectral exponent values that fall within the expected range of anomalous diffusion (see Supplementary Discussion 1).

**Figure 4:**
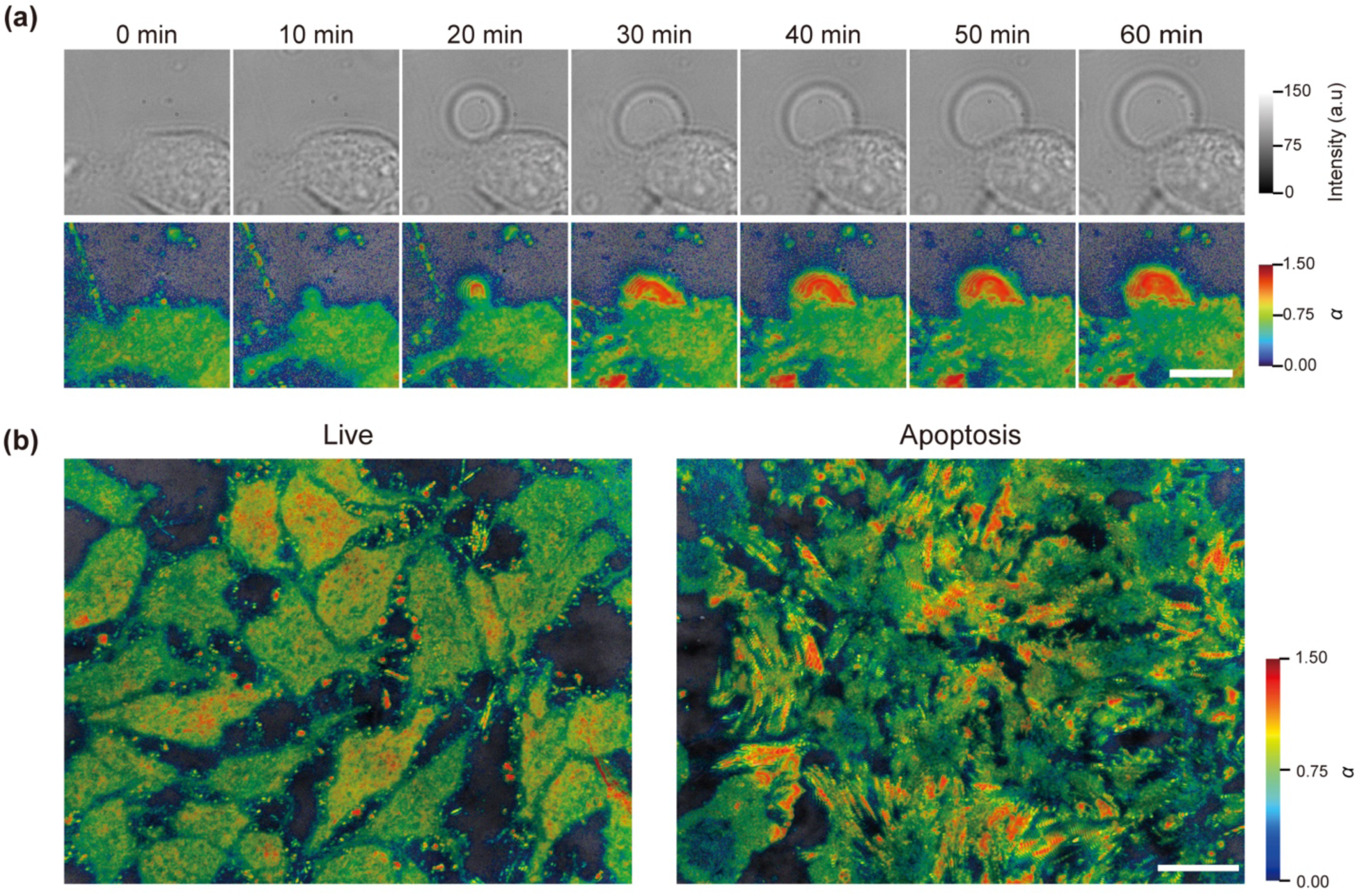
Visualization of spectral exponent changes in spiking HEK cells undergoing apoptosis. (a) Time-lapse brightfield images (top row) and corresponding 2D spectral exponent maps (bottom row) of spiking HEK cells over 1 hour following H₂O₂ treatment. Brightfield images showed progressive morphological changes, including blebbing, while 2D spectral exponent maps revealed the emergence of high *α* regions (red) near the cell periphery. Scale bar: 10 μm. (b) Comparison of the 2D spectral exponent maps in live and apoptotic spiking HEK cells, where apoptotic cells showed increased spatial heterogeneity of the spectral exponents, with the centers appearing blue and the peripheries red. Scale bar: 20 μm.

Fig. 4(b) compares 2D spectral exponent maps between live and apoptotic spiking HEK cells. In live cells, the cells predominantly displayed green-yellow hues (0.6 < *α* < 0.8), and some displayed slightly higher *α* values with yellow-orange hues (0.8 < *α* <1). For each live cell, the hues were relatively consistent throughout the whole cell body. In contrast, apoptotic cells primarily exhibited blue hues in the central regions (0.3 < *α* < 0.6) while featuring yellow-red hues (1< *α* < 1.5) in the peripheral regions, particularly at structures such as membrane protrusions and blebs. The 2D spectral exponent maps of apoptotic cells demonstrate greater spatial heterogeneity compared to those of live cells.

### 2.5 Cancerous cells differing in malignancy exhibit distinct spectral exponents

We further investigated whether the spectral exponent correlates with the aggressiveness of thyroid carcinoma subtypes. Papillary, follicular, and anaplastic thyroid carcinomas (PTC, FTC, and ATC) represent cancerous cells with increasing levels of invasiveness and malignancy. These subtypes originate from thyroid follicular epithelial cells, with PTC being the most common (∼80% of cases), FTC less frequent (10–20%), and ATC rare but highly aggressive (∼2%). Malignant thyroid cells are characterized by reduced cytoskeletal organization, fewer stress fibers, and decreased mechanical stiffness—up to 3- to 5-fold softer than normal thyroid cells (*36–38*).

Fig. 5(a) presents 2D spectral exponent maps of PTC, FTC, and ATC cells. PTC exhibited more extensive yellow to orange hues (*α* > 0.85), especially on the periphery. FTC cells displayed predominantly green to yellow hues (0.7 < *α* < 1.2). In contrast, ATC cells showed high *α* values at their leading/protruding edges with a dominance of green hues (0.4 < *α* < 0.9) at the center. These visual nuances are more apparent in the histograms of *α*-values (Fig. 5(b)). Both PTC and FTC cells, which share similarities as differentiated thyroid cancers, exhibited bimodal distributions, characterized by a primary peak at ∼0.93 and a secondary peak at lower *α* values. Notably, PTC cells showed a higher median α value (0.90) than FTC cells (0.82), while the dedifferentiated ATC cells exhibited the lowest median α value (0.67) and a distinct histogram profile likely due to a greater dissimilarity from the other two cell lines. Analysis of a larger sample pool (N=12) confirmed this trend: As plotted in Fig. 5(c), PTC, FTC, and ATC cells showed decreasing median α values (0.89, 0.79, and 0.71, respectively) correlated with their increasing malignancy, underscoring the potential of utilizing the 2D spectral exponent map as an intrinsic biomarker of cancer malignancy (*39*).

**Figure 5:**
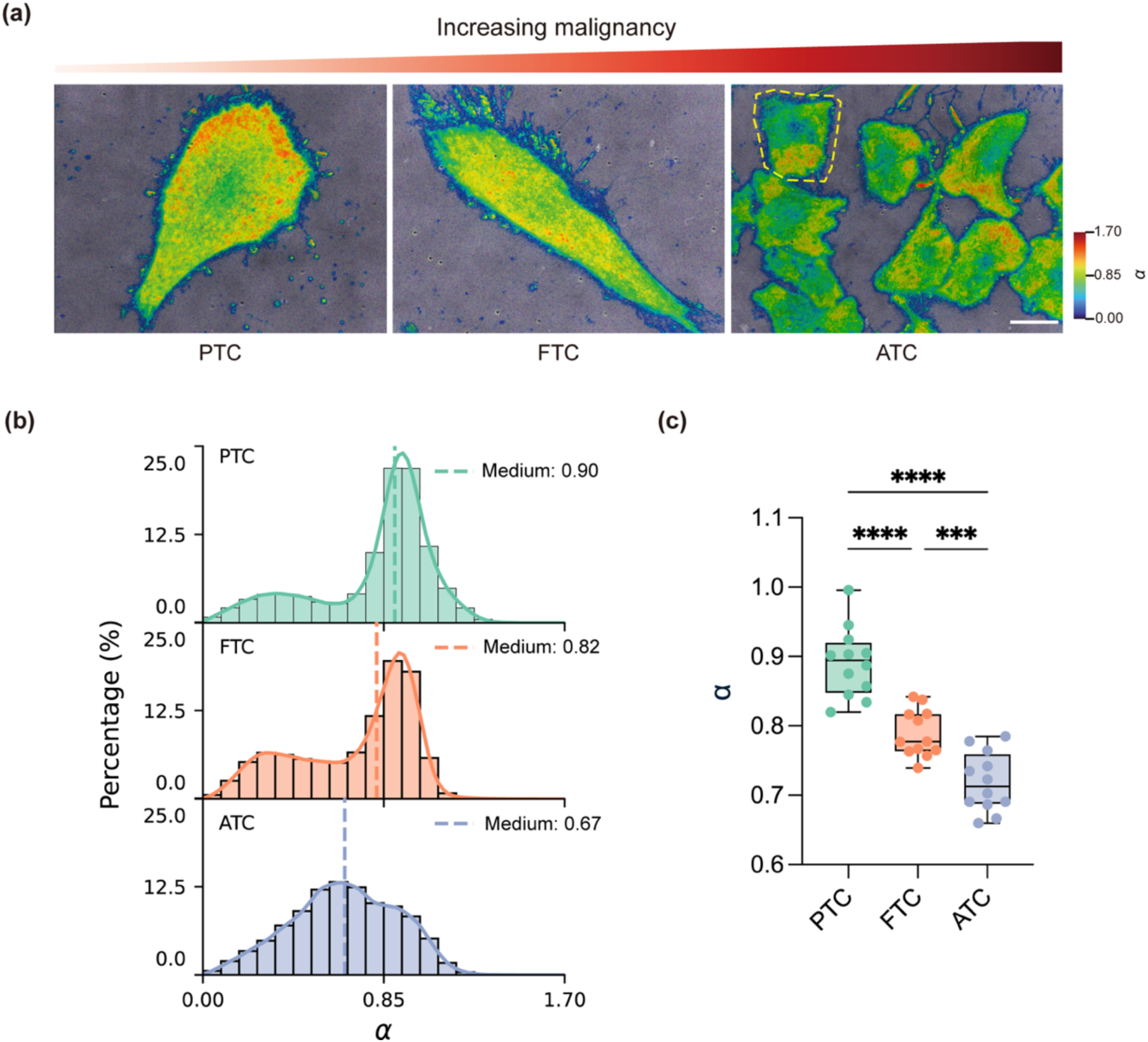
Correlation between the malignancy of thyroid carcinoma cancer cells and their spectral exponents. (a) 2D spectral exponent maps of PTC, FTC, and ATC cells. The yellow dash line in ATC indicates the single cell for plotting the histogram in (b). (b) Histograms of *α*-values extracted from the PTC, FTC and ATC cells in (a). (c) Comparison of the median *α*-values across three cancer cell lines. Pairwise comparisons were all statistically significant (Tukey’s test, ****: p < 0.0001 for PTC vs. FTC and PTC vs. ATC; ***: p < 0.001 for FTC vs. ATC), indicating a continuous decline in the median *α*-value with increasing malignancy. Sample size: N = 12 cells per group. Scale bar: 20 µm.

## 3. Discussion

In this study, we demonstrated power-law-based spectral exponent analysis to characterize subcellular dynamics in wide-field iSCAT imaging. By analyzing the temporal contrast signals in iSCAT and computing the PSD, we observed a power-law relationship over 30 - 1250 Hz that applied to most regions covered by live cells. Such a relationship was validated across diverse cell types. Mapping the spectral exponent of the power-law curve allows label-free imaging of live cells and offers insights into their physiological states.

Specifically, we demonstrated that the proposed method effectively captured dynamic transitions during mitosis (see *Results 2.3*), where cells in pro/metaphase exhibited high *α*-values, which may reflect the rapid restructuring and membrane remodeling that occur during chromatin condensation and spindle assembly (*40*). As cells re-entered interphase, *α*-values decreased, consistent with a return to a less active state and reduction of activities. In line with the spectral exponent changes observed during apoptosis (see *Results 2.4*) and across different malignancies (see *Results 2.5*), these findings suggest that the subcellular movements revealed in the power-law curve may be correlated with the structural organization or functional activities in live cells. Moreover, the internalized 200 nm AuNPs exhibited high α-values (Fig. 3(b)), which is perhaps related to their sub-diffusive motions in the cytoplasm or the relatively directional active transport that occurs when these nanoparticles are mechanically coupled to elastic or tensioned membranes (*41*, *42*). The notable difference in the spectral exponent of the internalized AuNPs compared to the intrinsic spectral exponent range of live cells allows for the direct identification of AuNPs in live cells. Such imaging capability may benefit the study of cellular uptake of nanoparticles for photothermal and photodynamic therapies, as well as for nanoparticle-based drug delivery.

The power-law relationship in the PSD is often associated with the diffusive thermal motion of scattering objects in solution (*43*, *44*). While a spectral exponent within the range of 1 < *α* < 2 aligns with the theoretical model of ensembled iSCAT signals of monodispersed scatterers undergoing sub-diffusion (see Supplementary Discussion 1), it is difficult to explain the spectral exponents smaller than 1 observed in some live cells. More efforts are needed to model the movements of scatterers in live cells and their implications for iSCAT signals to understand the mechanistic origin of the observed power-law relationship.

In conclusion, we have established spectral exponent analysis as a powerful tool to characterize stochastic fluctuations in iSCAT signals. This intrinsic biomarker effectively differentiates between different states and types of biological cells without relying on external markers. Spectral exponent analysis of iSCAT signals opens a broad avenue for applications in cancer diagnostics, stem cell differentiation, and live/dead cell assay in cell therapies. This technique may also be extrapolated to other interferometric imaging modalities, such as dynamic optical diffraction tomography (ODT) imaging and dynamic OCT imaging of thick tissues and organoids.

## 4. Materials and methods

### 4.1 iSCAT imaging setup

We constructed a custom wide-field iSCAT setup as depicted in Supplementary Fig. 4. A supercontinuum laser (SuperK FIANIUM FIU-6 OCT, NKT Photonics, A/S, Birkerød, Denmark) was employed as the illumination source instead of a conventional laser diode. The laser output was filtered using a combination of a long-pass dichroic mirror (DMLP950, Thorlabs), a 500 nm long pass filter (FELH0500, Thorlabs), a 700 nm short-pass filter (FES0700, Thorlabs) to achieve a wavelength range of 530–700 nm. The broadband illumination reduced the coherence length to ∼0.8 μm and ensured that the detected signals primarily originated from the interference between the reference beam reflected from the glass-water interface and the light scattered from the bottom section of the cells. The filtered beam was expanded from 1.3 mm to 8 mm in diameter using telescopes composed of lens pairs of 50 mm and 75 mm, and 75 mm and 300 mm. An achromatic lens (L1) with a focal length of 400 mm (AC508-400-AB, Thorlabs, USA) was used to focus the beam with the back focal plane of the microscope objective lens. We incorporated a low magnification oil-immersion objective lens (CFI Plan Fluor 20XC MI, Nikon Instruments Inc., NA = 0.75) to enlarge the field of view of the wide-field iSCAT system.

To separate the illumination and imaging paths, a polarizing beam splitter (PBS, CCM1-PBS251/M, Thorlabs, USA) was positioned after the first lens. A coupling mirror (M2) was mounted at a 45° angle to direct the illumination beam vertically toward the microscope objective, while a quarter waveplate (QWP, AQWP10M-580, Thorlabs, USA) was added after M2 to circularly polarize the illumination light before it reached the back aperture of the microscope objective. The microscope objective collected both the scattered light from the sample and the reflected light from the glass-water interface (the top surface of the coverslip), which became s-polarized after passing through the QWP the second time. The s-polarized light was then reflected by the PBS and focused onto a high-speed sCMOS camera (Phantom VEO 1310) using an imaging lens (L2, ACT508-1000-A, Thorlabs, USA). Additionally, a brightfield imaging path was incorporated to facilitate sample localization. An LED light positioned above the sample stage provided illumination for the brightfield imaging.

The wide-field iSCAT imaging setup had an overall magnification of 100 times, resulting in an effective pixel size of 0.18 µm/pixel. In our experiment, we captured each iSCAT video at 5000 frames per second with a camera resolution of 800 x 600 pixels, covering a field of view of 144 x 108 µm².

### 4.2 Cell culture for iSCAT imaging

#### a. Cell fixation

U2OS cells (ATCC® HTB-96™) were maintained in Dulbecco’s Modified Eagle Medium (DMEM) GlutaMAX™ (Gibco™ 10566016, Thermo Fisher Scientific) supplemented with 10% fetal bovine serum (FBS, Gibco™ A5256701, Thermo Fisher Scientific) and 1% penicillin-streptomycin (PS, Gibco™ 15140122, Thermo Fisher Scientific). Cells were incubated at 37 °C with 5% CO2 and plated on Poly-D-lysine (PDL, Gibco™ A3890401, Thermo Fisher Scientific)-coated 35-mm glass-bottom dishes (ibidi 81158) before imaging. Fixed cells were prepared by washing with warm phosphate-buffered saline (PBS, Gibco™ 10010023, Thermo Fisher Scientific), followed by fixation with 4% paraformaldehyde (Thermo Fisher Scientific J61899.AK) in PBS for 15 minutes. The cells were then washed three times with PBS.

#### b. Live and apoptotic cell imaging

Spiking HEK cells, originally developed by Adam Cohen’s group at Harvard University (*45*), were cultured in Dulbecco’s Modified Eagle Medium/Nutrient Mixture F-12 (DMEM/F12, Gibco™ 11320033, Thermo Fisher Scientific), supplemented with 10% FBS, 1% PS, 500 μg/mL geneticin (Gibco™ 10131035, Thermo Fisher Scientific), and 2 μg/mL puromycin (Gibco™ A1113803, Thermo Fisher Scientific). For live and apoptotic cell imaging, spiking HEK cells were plated on glass-bottom dishes (ibidi 81158) and cultured in DMEM/F12 supplemented with 10% FBS, 1% PS, 500 μg/mL geneticin, and 2 μg/mL puromycin for 24 hours before imaging. Before imaging, the cells were washed with DMEM/F12, no phenol red (Gibco™ 21041025, Thermo Fisher Scientific), and imaged in fresh DMEM/F12, no phenol red to capture live cell images. To induce apoptosis, 1.6 mM hydrogen peroxide (Sigma-Aldrich H1009) in DMEM/F12, no phenol red was added to the cells, followed by incubation at 37 °C with 5% CO_2_ for 30 minutes. The cells were subsequently imaged using iSCAT. For continuous monitoring, the images were taken every 10 minutes up to 60 minutes.

#### c. Mitosis imaging

HeLa cells were plated on glass-bottom dishes with 500-µm grids (ibidi 81168) and cultured in DMEM High Glucose (Gibco™ 11965092, Thermo Fisher Scientific) supplemented with 10% FBS and 1% PS for 24 hours before imaging. Cells exhibiting spherical morphology, indicative of prophase and metaphase stages, were identified under brightfield microscopy. iSCAT images were acquired at time points 0, 30, 45, and 100 minutes to monitor mitotic progression. Imaging was performed at 37 °C with 5% CO_2_ to maintain physiological mitotic conditions. A supercontinuum laser with a filter of 830 ± 15 nm was used to allow for longer coherent length into the rounded cells undergoing mitosis.

#### d. Cancer malignancy imaging

For cancer malignancy imaging, three thyroid cancer cell lines were analyzed: papillary thyroid carcinoma (PTC, K1), follicular thyroid carcinoma (FTC, RO82W1), and anaplastic thyroid carcinoma (ATC, SW1736). PTC and FTC cells were cultured in a complete medium composed of DMEM: F12:MCDB105 (Sigma-Aldrich 117-500) in a 2:1:1 ratio, supplemented with 10% FBS and 1% PS. ATC cells were cultured in RPMI 1640 (Gibco™ 11875-093, Thermo Fisher Scientific) supplemented with 10% FBS and 1% PS.

To induce cell cycle arrest, all three cell lines were serum-starved in serum-free media for 4 hours. Imaging was performed in a complete culture medium. Due to the different cell sizes, PTC and FTC cells were acquired individually in each field of view, while multiple ATC cells were captured in one field of view. For PTC and FTC imaging, the camera resolution was adjusted to 640 × 480 pixels to fit the cell sizes.

### 4.3 Data processing

#### a. Spectral exponent analysis of iSCAT signals

Each iSCAT recording was acquired at a frame rate of 5000 Hz. To reduce shot noise while preserving relevant dynamic information, two adjacent frames in the recorded iSCAT videos were binned, resulting in a reduced frame rate of 2500 Hz. This frame rate was selected to capture dynamics up to 1250 Hz to ensure an adequate spectral range. The total duration of each video was ∼5 seconds. Variation in laser intensity was corrected by multiplying a calibration coefficient for each frame so that the background intensity remained consistent throughout the recording. The contrast signal *I*(*t*) was calculated from the raw iSCAT signal *I*_*d*_(*t*) following the equation:

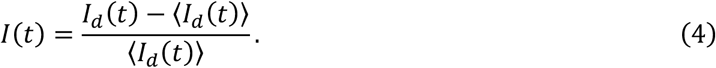

To analyze temporal fluctuations at each pixel, we computed the PSD using Welch’s method. We applied a window size of 300 frames to reduce the influence of noise on the PSD estimation. The PSD, denoted as *S*(*f*), is defined as the square of the magnitude of the Fourier transform of the signal *I*(*t*):

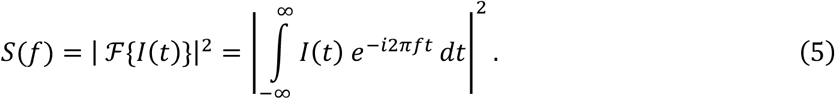

Subsequently, the resulting PSD for each pixel was fitted to an inverse-power-law model (Eq. 3), with all parameters constrained to be greater than zero during the fitting process. To minimize potential biases due to mechanical instabilities, the lowest frequencies (i.e., the bottom ∼3% of the spectrum, corresponding to <30 Hz) in the PSD were excluded from the analysis.

### b. Evaluating fitting performance

To evaluate how the inverse-power-law model fits experimental data, we employed the coefficient of determination (*R*²) as a metric for quantifying the goodness of fit. The *R*² value measures the proportion of variance in the observed PSD, *S*(*f*), that can be explained by the model prediction, *Ŝ*(*f*). A higher *R*² value, closer to 1, signifies a better fit, while a value near 0 suggests that the model fails to explain the observed variability. The *R*² value was calculated as:

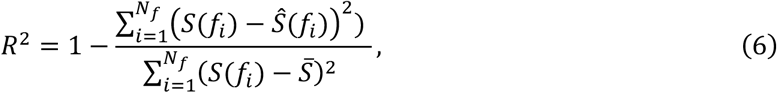

where *S̄* represents the mean of the observed PSD values across the entire frequency range. The denominator quantifies the total variance in the observed data, while the numerator represents the residual variance not explained by the model.

### c. Constructing the 2D spectral exponent map

We introduced a color-coded 2D spectral exponent map to better visualize the spatial distribution of the fitted parameters in the inverse-power-law model. Considering an HSV image consisting of the hue, intensity, and saturation channels, the *α*-values were mapped to the hue channel using the Turbo colormap provided by MATLAB (MathWorks), instead of the standard HSV hue scale. This substitution was made to enhance perceptual contrast and improve visual differentiation across spatial regions. The intensity value (*IV*) was obtained by calculating the predicted power spectral density *Ŝ*(*f*) at the lowest frequency (*f* = 30 Hz), which provided a consistent and interpretable reference point across all pixels and datasets, as the intensity value was evaluated at the same frequency regardless of pixel location. To compress the dynamic range and improve contrast, the *IV* values were log-transformed. The saturation channel was adjusted by *R*^2^ to ensure that regions with a stronger power-law fit appear more vivid, while areas with lower *R*^2^ remain less prominent.

Min-max normalization was applied to *α*, *IV*, and *S*, rescaling each parameter to a [0, 1] range based on their minimum and maximum values:

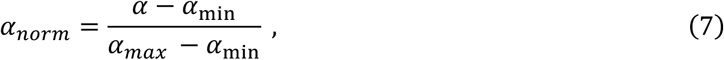

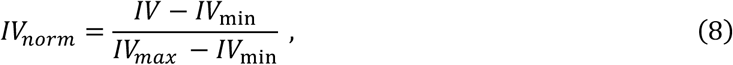

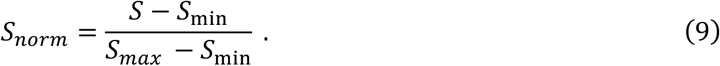

Finally, the HSV image was converted into an RGB image, allowing for effective visualization of the spatial variations in the power-law parameters.

### d. Obtaining the mean frequency map for comparison with the 2D spectral exponent map

To calculate the mean frequency map, the PSD *S*(*f*) was first normalized by the total power,

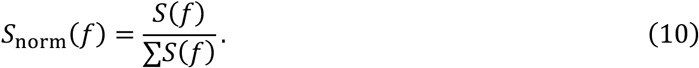

The mean frequency for each pixel was calculated as a single parameter to describe the dominant dynamics within the pixel’s temporal signal following the equation,

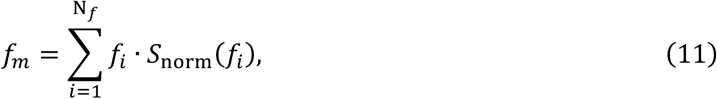

where *N*_*f*_ is the total number of frequency points in the PSD.

### e. Statistical analysis

To generate histograms and determine median α values in a large sample pool, individual cell areas were segmented using MATLAB (MathWorks). Segmentation was performed on the mean frequency map, where cell regions were readily distinguishable from the background. Each cell was individually cropped, and pixels corresponding to the cell area were isolated by applying a threshold of <300 Hz to the mean frequency map to exclude pixels that typically correspond to background regions. The resulting cell masks were saved for the regions of interest (ROIs) in subsequent spectral exponent analyses. Statistical analyses, including boxplot generation, were conducted using Prism 10 (GraphPad Software). Two-tailed unpaired *t*-tests were performed to study the spectral exponents in live/fixed cells. The ordinary one-way ANOVA (with Tukey’s multiple comparisons test) was used to study the median spectral exponents in cancerous cells across various malignancies.

## Acknowledgements

We thank Ms. Xinjuan Koh for assisting with culturing spiking HEK cells, and Prof. Wenting Zhao’s group for sharing the U2OS and thyroid cancer cell lines. This work was funded by grants from National Research Foundation Singapore (NRF-NRFF14-2022-0005, T.L), the Startup Grant from Nanyang Technological University (T.L.), the Ministry of Education, Singapore under its AcRF Tier 2 Grant (MOE-T2EP30124-0010, T.L.) and AcRF Tier 1 Grant (RS19/20, T.L.; RG28/21, T.L.).

## Author contributions

T.L. conceived the idea. C.L.A. built the iSCAT setup. C.L.A., H.Y., and G.K.L. conducted the experiments. C.L.A., T.L., and H.Y. analyzed the data. C.L.A. conducted theoretical analysis. C.L.A., H.Y. and T.L. drafted the manuscript. C.L.A., H.Y., H.L., and T.L. discussed and revised the manuscript. All work was supervised by T.L.

