## Supplementary Document for "Label-Free Mapping of Subcellular Dynamics using Wide-field Interferometric Scattering Microscopy and Spectral Exponent Analysis"

**This supplementary information includes:**

Supplementary Discussion:

1. Theoretical analysis of the power spectral density (PSD) of ensembled iSCAT signals in monodispersed scatterers undergoing Brownian motion
2. Comparison of power-law and piecewise functions for spectral exponent analysis
3. Optomechanical stability assessment
4. Dynamics of live and fixed cells revealed by the 2D spectral exponent maps

Supplementary Figures 1 to 4

**Supplementary Discussion 1. Theoretical analysis of the power spectral density (PSD) of ensembled iSCAT signals in monodispersed scatterers undergoing Brownian motion**

**a) The ensembled iSCAT constrast signal**

According to Eq. (1) in the main text and previous studies in live cell imaging using iSCAT (*1*, *2*), the second and fourth terms in Eq. (1) can be omitted due to $|r| \gg |\kappa_n|$. In this case, the raw iSCAT signal can be expressed as,

$$I_d(t) \approx |E_{\text{inc}}|^2 \left[ |r|^2 + 2 \sum_n |r||\kappa_n| \cos[2kz_n(t) + \theta_n] \right], \quad \text{(S1)}$$

where $r$ denotes the reflection coefficient of the glass-water interface, $k$ represents the center wave number, $z_n(t)$ represents the optical path difference between the reference light and the light scattered from the $n$-th scatterer, $\kappa_n$ denotes the complex polarizability of the $n$-th scatterer, and $\theta_n = \arg \kappa_n - \arg r$, which accounts for the phase difference between $r$ and $\kappa_n$.

Due to the strong intensity of the illumination reference beam and the stochastic nature of the second term in Eq. (S1), the illumination reference beam $I_r = |E_{\text{inc}} r|^2$can be estimated by averaging $I_d(t)$ over time (*1*). The contrast $I(t)$ of the raw iSCAT signal $I_d(t)$ can be obtained by,

$$I(t) = \frac{I_d(t) - I_r}{I_r}. \quad \text{(S2)}$$

Because $|r| \gg |\kappa|$, $I(t)$ can be approximated to,

$$I(t) \approx \frac{2}{|r|} \sum_n |\kappa_n| \cos(2kz_n(t) + \theta_n). \quad \text{(S3)}$$

For simplicity, here we use $\phi_n'(t)$ to represent $2kz_n(t) + \theta_n$. We can thus obtain,

$$I(t) \approx \frac{2}{|r|} \sum_n |\kappa_n| \cos\big(\phi_n'(t)\big). \quad \text{(S4)}$$

**b) Autocorrelation of the iSCAT contrast signal**

To relate the fitted power spectral density (PSD) to the stochastic thermal motion of scatterers, we consider a monodispersed solution in which all the scatterers share the same motion statistics. To analyze their ensembled iSCAT signal, we define the autocorrelation of the iSCAT contrast signal $I(t)$ as,

$$\Gamma(\tau) = \langle I(t) I(t+\tau) \rangle. \quad \text{(S5)}$$

Here, $I(t)$ represents the temporal contrast signal at a fixed pixel location in the iSCAT video, and $I(t+\tau)$ is the same signal delayed by a lag time $\tau$. The angle brackets $\langle \cdot \rangle$ denote a temporal average over time $t$.

By substituting $I(t)$ from Eq. (S4) into Eq. (S5), we obtain,

$$\Gamma(\tau) = \left\langle \left( \frac{2}{|r|} \sum_n |\kappa_n| \cos\big(\phi_n'(t)\big) \right) \left( \frac{2}{|r|} \sum_m |\kappa_m| \cos\big(\phi_m'(t+\tau)\big) \right) \right\rangle \ . \tag{S6}$$

Upon expanding the product and assuming that cross-terms (for $m \neq n$) average to zero due to random, uncorrelated phases, the autocorrelation can be simplified to:

$$\Gamma(\tau) = \frac{4}{|r|^2} \sum_n |\kappa_n|^2 \langle \cos \phi_n' \,(t) \cdot \cos \phi_n' \,(t+\tau) \rangle \, . \tag{S7}$$

Using the trigonometric identity:

$$\cos A \cos B = \frac{1}{2} [\cos(A - B) + \cos(A + B)], \tag{S8}$$

and noting that the second term of Eq. (S8) averages to zero, the autocorrelation term becomes,

$$\Gamma(\tau) = \frac{2}{|r|^2} \sum_n |\kappa_n|^2 \langle \cos\big(\Delta\phi_n'(t)\big) \rangle \, , \tag{S9}$$

where $\Delta\phi_n'(t, \tau) = \phi_n'(t) - \phi_n'(t + \tau)$.

For monodispersed scatterers undergoing classical diffusion, the axial displacement, denoted as $\Delta z_n$, follows a Gaussian distribution, and its mean square displacement (MSD) grows as:

$$\langle |\Delta z_n|^2 \rangle = 2 D_z \tau, \tag{S10}$$

where $D_z$ is the diffusion coefficient along the z-axis. Consequently, the mean square phase shift is given by:

$$\langle |\Delta\phi_n'|^2 \rangle = 4k^2 \langle |\Delta z_n|^2 \rangle = 8k^2 D_z \tau. \tag{S11}$$

Assuming $\Delta\phi_n'$ is normally distributed, we can use the identity $\langle \cos(\Delta\phi_n') \rangle = \exp\left(-\frac{\langle |\Delta\phi_n'|^2 \rangle}{2}\right)$, and substitute the corresponding term in Eq. (S9), which leads to:

$$\Gamma(\tau) = \Gamma_0 e^{-4k^2 D_z \tau}, \tag{S12}$$

where $\Gamma_0 = \frac{2}{|r|^2} \sum_n |\kappa_n|^2$ is the autocorrelation amplitude at zero lag time. This exponential decay is a feature of the ensembled iSCAT signal that originates in monodispersed scatterers undergoing diffusive motion.

**c) Obtaining the PSD of the iSCAT contrast signal via Wiener-Khinchin theorem**

According to the Wiener–Khinchin theorem, the PSD is the Fourier transform of the autocorrelation function (*3*). Therefore, the PSD of the temporal contrast signal $I(t)$ can be expressed as:

$$S(f) = \int_{-\infty}^{\infty} \Gamma(\tau) \, e^{-i2\pi f \tau} \, d\tau. \tag{S13}$$

Substituting Eq. S12 into Eq. S13 yields a Lorentzian PSD,

$$S(f) = \frac{2\Gamma_0 k^2 D_z}{(2k^2 D_z)^2 + (\pi f)^2} \,. \tag{S14}$$

This $S(f)$ derived from the autocorrelation function of classical diffusion (Brownian motion), cannot be represented as a simple power law across the entire frequency range due to its Lorentzian shape. However, in certain limits or ranges of $f$, we can approximate a power-law behavior, where, for very high frequencies when $f \gg k^2 D_z$, the $(\pi f)^2$ term dominates the denominator, and the PSD can be approximated to,

$$S(f) \approx \frac{2\Gamma_0 k^2 D_z}{(\pi f)^2}. \tag{S15}$$

In this regime, the PSD exhibits a $f^{-2}$ dependency, which is a form of power-law behavior with $\alpha = 2$. This reflects a typical decay rate for systems dominated by Brownian motion or classical diffusion. At low frequencies when $f \ll k^2 D_z$, the constant term $(2k^2 D_z)^2$ in the denominator is more significant, and the PSD does not simplify to a power law form but rather approaches a constant value determined by $\Gamma_0$, reflecting static contributions from the scatterers.

If the monodispersed scatterers undergo anomalous diffusion instead of Brownian motion, the PSD in Eq. (S15) can be generalized to a power-law equation:

$$S(f) \approx \beta f^{-\alpha}. \tag{S16}$$

Here, sub-diffusion ($1 < \alpha < 2$) is common in environments with obstacles or mechanical constraints, such as those imposed by the cytoskeleton and macromolecular crowding, whereas super-diffusion ($\alpha > 2$) may indicate active movements driven by molecular motors or directed transport.

**d) Implications on the spectral exponent analysis of the ensembled iSCAT signal in live cells**

The analysis presented above introduces a possible mechanism that highlights the power-law relationship observed in the PSD of the ensembled iSCAT signal from monodispersed scatterers experiencing stochastic thermal motion. Meanwhile, it does not cover the regime of $0 < \alpha < 1$ we observed in the experimental results. In fact, modeling the ensembled iSCAT signal in live cells is considerably more complicated due to the inherent heterogeneity of the scatterers, which include membranes, coacervates, and individual proteins, as well as their complex environment subject to compartmentalization, clustering, and crowding. It remains a challenge to provide a plausible mechanistic explanation for the power-law relationship we have observed in live cells. Nevertheless, this work demonstrates how such an empirical relationship correlates with cellular activities, such as mitosis, apoptosis, and malignancy. Further studies are necessary to shed light on the mechanistic insight and elucidate whether it could be related to a similar mechanism as introduced in Eqs. (S1) - (S16).

**Supplementary Discussion 2. Comparison of piecewise and power-law functions for spectral exponent analysis**

In our fitting function, we adopted a piecewise approach to determine the noise floor of each pixel:

$$\hat{S}(f) = \begin{cases} \beta f^{-\alpha}, & f < f_c \\ \eta, & f \geq f_c \end{cases}. \tag{S17}$$

Here, $f_c$ is the transition frequency (critical frequency where the power-law trend discontinues), and $\eta$ represents the amplitude of the noise floor. This formulation allows us to restrict the fitting operation to the frequency range where actual power-law scaling is observed.

For comparison, we also considered a global power-law model applied to the full spectral range,

$$\hat{S}(f) = \beta f^{-\alpha} + \eta. \tag{S18}$$

In this model, the noise floor term $\eta$ is incorporated across all frequencies. In practice, both Eq. (S17) and Eq. (S18) achieved good fitting performance when the signal-to-noise ratio was high (see the fitting results at the 1st and 2nd pixels in Supplementary Fig. 1). However, a severe problem with fitting via Eq. (S18) was the overfitting problem when the PSD consisted of primarily the noise floor (see the 3rd pixel in Supplementary Fig. 1). In this case, the problem became ill-posed as changing either $\eta$ or $\beta$ in Eq. (S18) while making $\alpha$ close to 0 would constitute a valid solution, which resulted in the high fluctuation in the $\beta$ value estimated for the image background (Supplementary Fig. 1b). Although this problem can be mitigated by adding a regularization term in the optimization loss function, the additional regularization term may sometimes affect the fitting results and requires specific attention of the end users. Therefore, we chose Eq. (S17) for the spectral exponent analysis in this work.

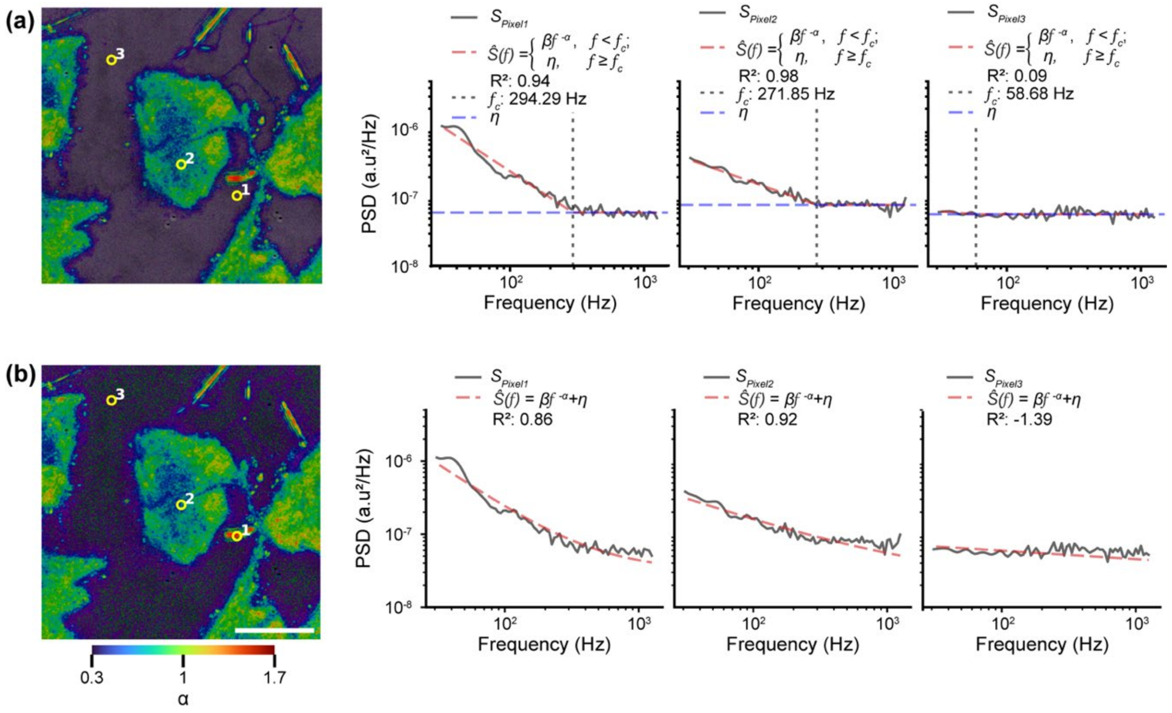


**Supplementary Fig. 1**. Comparison of 2D spectral exponent maps generated from (a) piecewise model (Eq. S17) and (b) global power-law model (Eq. S18). PSDs from pixels labeled 1–3 were fitted using both methods, and the global power-law model shows higher fluctuations in background regions. Scale bar: 20 µm.

### Supplementary Discussion 3. Optomechanical stability assessment

To evaluate the optomechanical stability of our imaging system, we tracked the position of a single, immobilized 200 nm gold nanoparticle (AuNP) on a clean coverslip over time. The relatively large size of the AuNP provided high interferometric contrast, which minimized localization errors and enabled precise tracking. The measurement was performed over a timescale of 8 seconds at a frame rate of 5 kHz.

A representative iSCAT image of the immobilized AuNP is shown in Supplementary Fig. 2(a), alongside its corresponding $f_m$ map in panel (b) and 2D spectral exponent map in panel (c). The interferometric point spread function (iPSF) of the particle (Supplementary Fig. 3(d)) displays a full width at half maximum (FWHM) of 472.6 nm, which closely matches the theoretical diffraction-limited value (~470 nm, for illumination wavelength of 580 nm).

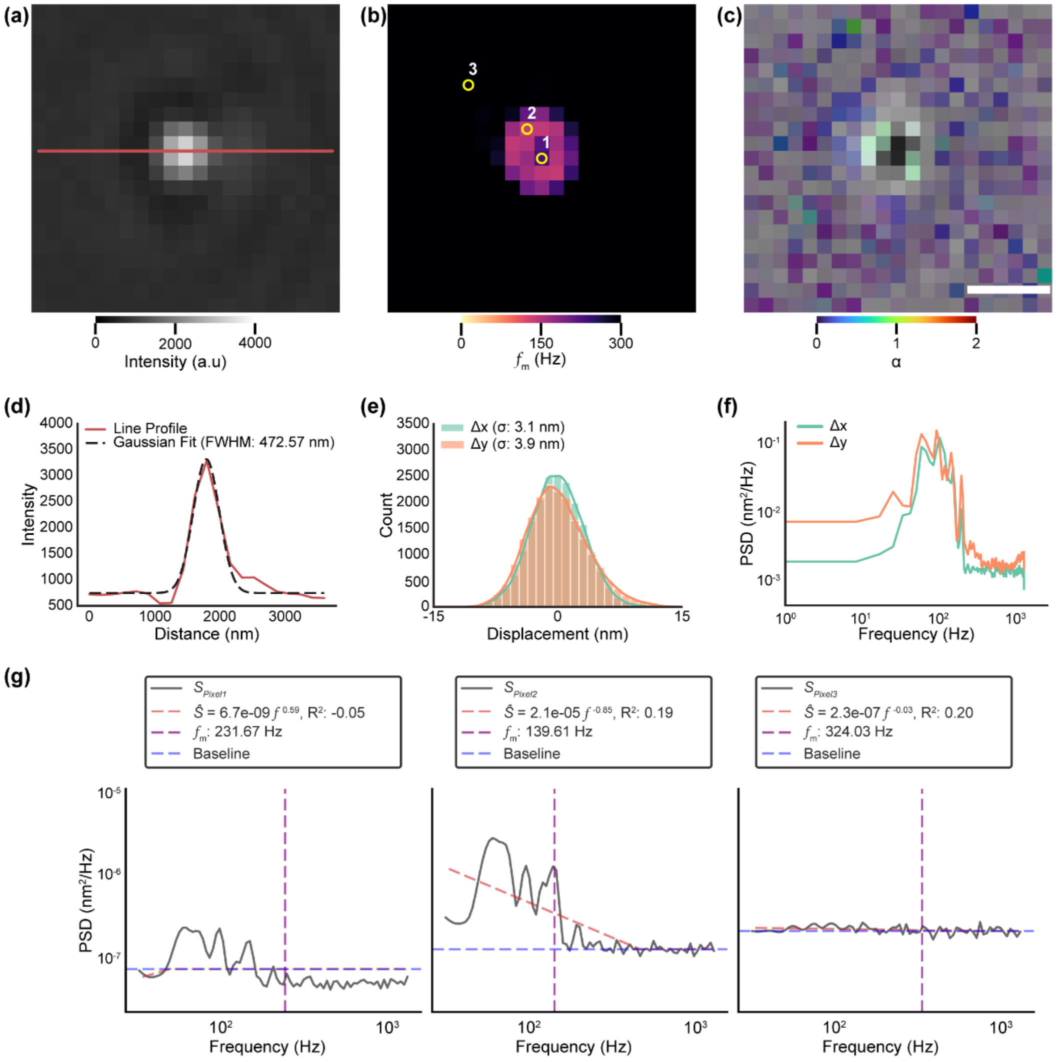


**Supplementary Fig. 2**. Optomechanical stability characterization of the iSCAT setup. (a) iSCAT image, (b) $f_m$-map and (c) 2D spectral exponent map of an immobilized 200 nm gold nanoparticle on a clean coverslip, captured at 5 kHz. (d) Line profile of an interferometric point-spread function (iPSF) of the single AuNP. The full width half max (FWHM) of the line profile is 472.6 nm. (e) Histogram of x and y displacements. (f) Fourier transform of $\Delta x$ and $\Delta y$ and (g) PSDs of pixels labeled 1, 2 and 3 in (b), fitted to Eq. S17. Scalebar: 5 μm.

Particle localization in both x and y directions was performed using the Trackpy algorithm. The histograms of positional displacements, as shown in Supplementary Fig. 2(e), reveal sub-nanometer fluctuations in both directions, with a root-mean-square (RMS) amplitude of 2.84 nm in the x-direction and 3.25 nm in the y-direction. Further vibration analysis reveals that the dominant motion occurs at frequencies between 100–140 Hz (Supplementary Fig. 2(f)). These unwanted mechanical instabilities introduce broadband background signals that are captured by the $f_m$-map, as the mean frequency calculation integrates the contributions from all frequency components in the PSD and biases the result. In contrast, the power-law fitting method is more resilient to such mechanical artifacts. Because the power-law fitting focuses on scale-free temporal dynamics, it does not converge for immobilized particles lacking true biological fluctuations, resulting in low $R^2$ values (Supplementary Fig. S3(g)) and effectively suppressing vibrational noise. This result highlights the advantage of the power-law analysis in isolating biologically relevant activities while minimizing interference from mechanical instabilities.

**Supplementary Discussion 4. Dynamics of live and fixed cells revealed by the 2D spectral exponent maps**

To assess the utility of the proposed method, we compared dynamic signal profiles between live and fixed U2OS cells. This comparison served as a reference baseline for interpreting signal fluctuations associated with cellular activity versus non-active states. The 2D spectral exponent map (Supplementary Fig. 3(a)(ii)) of live U2OS cells highlights the spatial distribution of the cell dynamics. Similar to the $f_m$-map in Supplementary Fig. 3(a)(i), we can clearly differentiate cell morphology without the use of exogenous markers.

The dynamics of U2OS cells underwent substantial changes upon fixation, as observed in the $f_m$-map (Supplementary Fig. 3(b)(i)) and the 2D spectral exponent map (Supplementary Fig. 3(b)(ii)). Compared to live cells, the fixed U2OS cells exhibited a reduction in dynamic heterogeneity and an overall increase in the mean frequency. The fluctuation signal was mainly shown in the cytosol region, where the mean frequency value was higher than that in live cells. Notably, the nucleus region exhibited predominantly white noise ($f_m$ close to 300 Hz), confirming the absence of subcellular dynamics. In line with the observation in the $f_m$-map, the colors in the 2D spectral exponent map changed from green to blue, indicating a drop in the spectral exponent and a flatter power-law curve. To assess these observations in a statistically meaningful manner, we compared the $\alpha$-distributions between live and fixed cells in 12 cells (Supplementary Fig. 3(c)). Live U2OS cells exhibited a higher median $\alpha$-value (1.02), whereas fixed cells showed lower α-values (0.46).

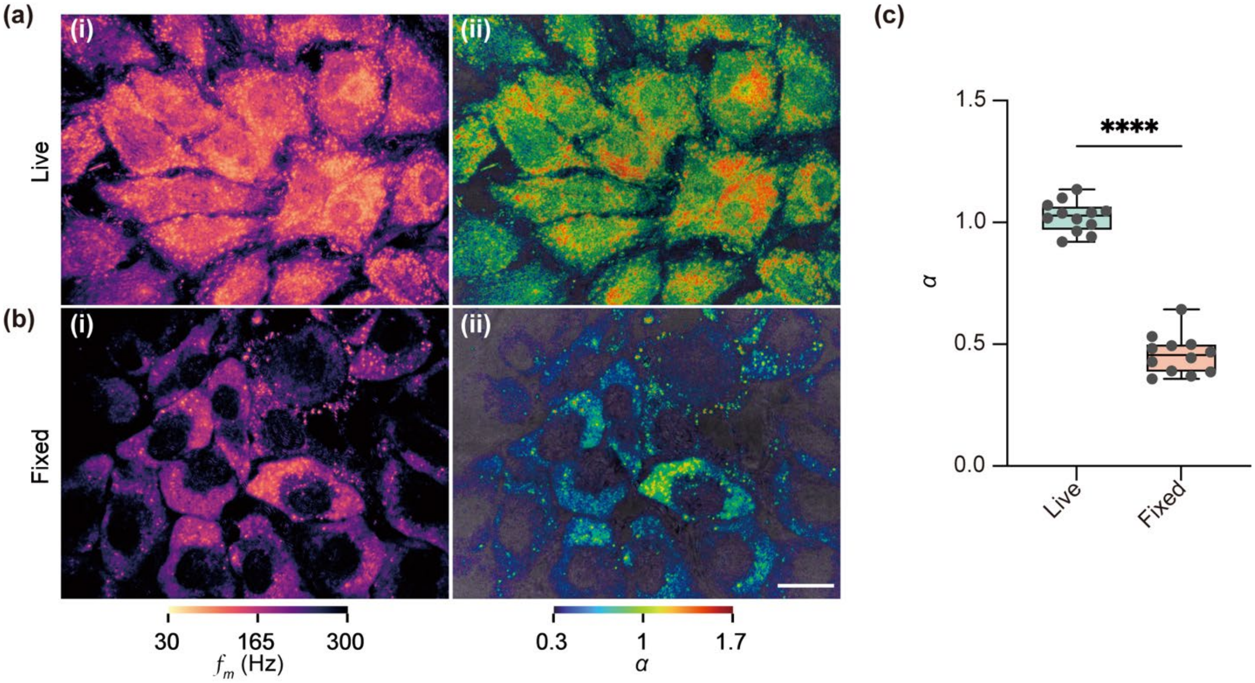


**Supplementary Fig. 3**. Comparison of the $f_m$-map and 2D spectral exponent map in imaging live and fixed U2OS cells. (a) $f_m$-maps and (b) 2D spectral exponent map visualizing the spatial distribution of the mean frequency and spectral exponent $\alpha$ in live and fixed U2OS cells. (c) Boxplot of the medium α-values comparing live and fixed U2OS cells. Each box represents the interquartile range, with the median shown as the central line; whiskers denote the full data range. Live cells exhibit significantly higher $\alpha$-values compared to fixed cells. The difference was statistically significant (****: $p < 0.0001$, two-tailed unpaired t-test; $t(22) = 19.10$, $R^2 = 0.9431$). Sample size: n = 12 cells per group. Scale bar: 20 $\mu$m.

### Experimental Setup

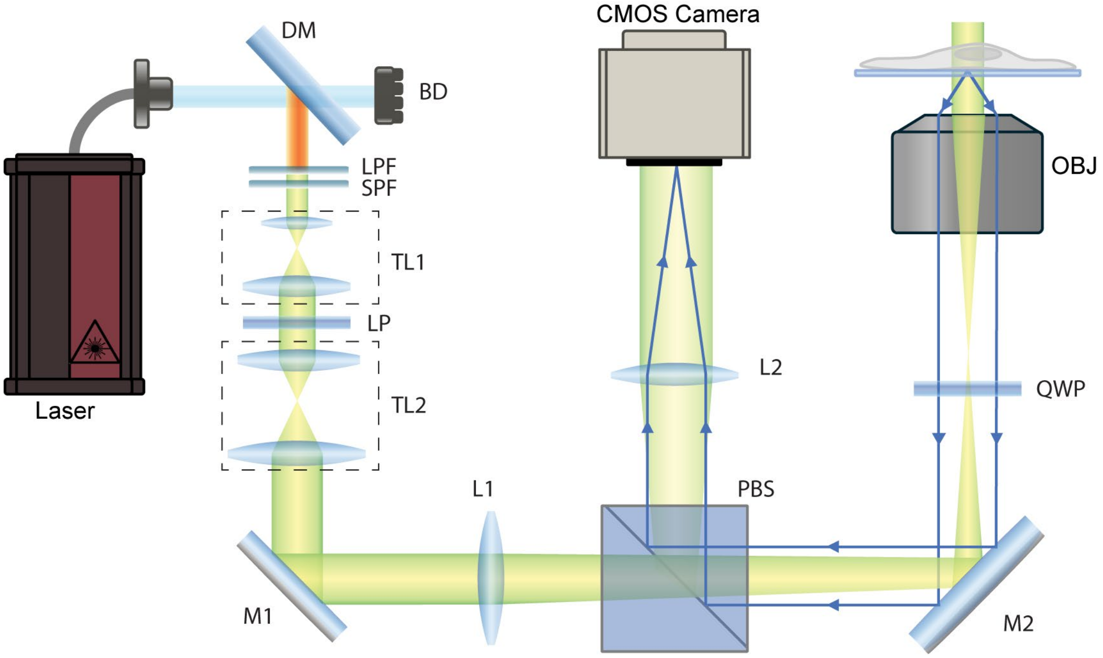


**Supplementary Fig. 4**. Schematic diagram of iSCAT experimental setup. OBJ: Objective lens, L1 and L2: Achromatic lens, M1 and M2: Mirrors, LPF: Long-pass filter, SPF: Short-pass filter, LP: Linear polarizer, QWP: Quarter-wave plate; PBS: Polarizing beam splitter, DM: Long-pass dichroic mirror, BD: Beam dump; TL1: Beam expander composed of 50 mm and 75 mm and TL2: Beam expander composed of 75 mm and 300 mm lens pairs, used to expand the beam from ~1.3 mm to ~8 mm diameter.
